# Screening of Cultivars for Tissue Culture Response and Establishment of Genetic Transformation in a High-yielding and Disease-resistant Cultivar of *Theobroma cacao*

**DOI:** 10.1101/2020.10.07.327486

**Authors:** Jesse Jones, Elaine Zhang, Dominick Tucker, Daniel Rietz, Doug Dahlbeck, Michael Gomez, Claudia Garcia, Jean-Philippe Marelli, Donald Livingstone, Ray Schnell, Brian Staskawicz, Myeong-Je Cho

## Abstract

A highly efficient transformation protocol is a prerequisite to developing genetically modified and genome-edited crops. A tissue culture system spanning the initiation of floral material to the regeneration of plantlets into soil has been tested and improved in cacao. Fourteen cultivars were screened for their tissue culture response and transfer DNA (T-DNA) delivery efficiency via *Agrobacterium*. These key factors were used to determine the genetic transformability of various cultivars. The high-yielding, disease-resistant cultivar INIAPG-038 was selected for stable transformation and the method was further optimized. Multiple transgenic events were produced using two vectors containing both yellow fluorescent protein and neomycin phosphotransferase II genes. A two-fold strategy to improve both T-DNA delivery and secondary somatic embryogenesis rates was conducted to improve overall transformation frequency. The use of *Agrobacterium* strain AGL1 and cotyledon tissue derived from immature somatic embryos ranging in size between 4-10 mm resulted in the highest T-DNA delivery efficiency. Furthermore, the use of higher concentrations of basal salts and cupric sulfate in secondary callus growth medium increased the percentage of explants producing greater than ten embryos by 504% and 443%, respectively. Consequently, an optimal combination of all these components resulted in a successful transformation of INIAPG-038 with 3.7% frequency at the T_0_ plant level. Grafting transgenic scions with undeveloped roots to wild-type seedlings with strong, healthy roots helped make plantlets survive and facilitated quick transplantation to the soil. The present methods can be applied to improve tissue culture response and transformation frequency in other cacao cultivars.

**Key message:** Tissue culture and genetic transformation methods for a high-yielding, disease-resistant cultivar of *Theobroma cacao* were established while factors affecting T-DNA delivery and somatic embryogenesis were identified.

## Introduction

*Theobroma cacao* L. (cacao) is an economically important crop grown predominantly in Africa, although the species was originally domesticated in South America and significant production still remains there. Despite cacao beans being a commodity in a multibillion-dollar industry, there have been severe losses from disease to both yields and trees (Fister et al. 2018; Marelli et al. 2019). Obstacles such as long juvenile period, issues with heterozygous and heterogeneous populations (Wickramasuriya and Dunwell 2018), long-life cycle and funding constraints on conventional cacao breeding programs (Maximova et al. 2003) make desired traits like high yield, disease resistance and bean quality difficult to combine into a single cultivar. However, both genome editing and transgenic approaches can be used to improve and greatly accelerate the pace of trait development.

The first stable transformation to regenerate cacao plants was reported by Maximova et al. (2003) and since then there have been other successful reports of the transformation of TcChil1 (Maximova et al. 2006) and TcLEC2 (Shires et al. 2017) into cacao. The introduction of BABY BOOM transcription factor into cacao significantly increased transformation frequency, but it caused a regeneration issue with an abnormal phenotype (Florez et al. 2015). These previously mentioned experiments were performed on the amenable cultivar PSU SCA-6. Previous to this study, PSU SCA-6 was the only cultivar that transgenic plants have been regenerated from. Although PSU SCA-6 is not identical to SCA-6, the original SCA-6 was collected from the wild in the Peruvian Amazon (Zhang et al. 2011) and has some natural, broad-spectrum fungal resistance (Pokou et al. 2019). However, this cultivar is not highly productive (Wahyu et al. 2009). For genetic transformation to be used most effectively the ability to transform many cacao cultivars, including those less-amenable to transformation, must be established. Ideally, a single genotype-independent transformation method would be developed, but similar to somatic embryogenesis which has had broad success across the plant kingdom, there still exists a high degree of genotype-to-genotype variation and so protocol customization is necessary (Garcia et al. 2016). Optimization of the basal salts, duration of hormone treatments, explant quality, and explant type are effective ways to improve the culturing response of recalcitrant cultivars.

Several explant types are used in cacao tissue culture and the transformation process, which include petals and staminodes from immature flower pods, cotyledon tissue from immature somatic embryos, and mature somatic embryos. Petals and staminodes are capable of primary somatic embryogenesis (Maximova et al. 2002) and are the starting point for all subsequent processes. Cotyledon tissue derived from somatic embryos is an alternative tissue type, efficient at secondary, tertiary and quaternary somatic embryogenic processes (Maximova et al. 2003). Mature somatic embryos undergo germination and are able to regenerate into plantlets. So far, all previous transformation methods (Maximova et al. 2003, 2006; Florez et al. 2015; Shires et al. 2017) used an embryogenic approach utilizing cotyledon tissue from somatic embryos. The advantages to this approach include previously described methodologies, the predominant single-cell origin of embryos (Maximova et al. 2002), and the high number of embryos cotyledon tissue are capable of producing. Alternatively, an organogenic approach was not thoroughly explored because of the lack of previously described methods in cacao, the apparent difficulty of establishing a new tissue culture system, and the heterogeneous nature of cacao seed which would result in the loss of the original cultivar if used as an organogenic donor material.

In this study we report a successful *Agrobacterium*-mediated transformation and regeneration protocol for INIAPG-038, using improved somatic embryogenesis and *Agrobacterium* infection. INIAPG-038 is a breeding line developed by Mars Wrigley, Incorporated in collaboration with USDA and INIAP. It is a highly productive cultivar which has tolerance to fungal diseases such as frosty pod disease (Costa et al. 2012; Meinhardt et al. 2014) and witches broom disease (Mondego et al. 2008). We also demonstrate a method for evaluating how amenable a cultivar is to tissue culture and transformation as well as provide data on the cultivars we screened. Outside of genetic transformation, improvement of somatic embryogenesis is useful for micropropagation.

## Materials and Methods

### Plant material

Fourteen cacao cultivars, SCA-6, CCN-51, INIAPG-038, INIAPT-374, INIAPT-072, INIAPG-152, INIAPM-226, INIAPM-053, INIAPM-196, INIAPG-029, INIAPT-484, GS-36, INIAPG-069 and Matina 1-6, were used for tissue culture response experiments. Flowers were shipped from USDA ARS Subtropical Horticulture Research Station in Miami, Florida. Flowers were harvested from the field and greenhouse in the morning then shipped overnight to the Innovative Genomics Institute (IGI) Plant Genomics and Transformation Facility, Berkeley, California, USA. Cultivars were primarily chosen based on their consistent availability of flowers.

### Initiation of primary embryogenesis using flower material and improvements

Immature flowers were used to initiate primary somatic embryos according to the previous protocols (Maximova et al. 2012; Garcia et al. 2016) with some modifications and the addition of various shipping methods. Explants of 14 cultivars were observed after 3-4 months of culturing to determine the rate at which both staminode and petal explants formed embryos. Between 250 and 2500 staminodes and petals were used for each cultivar across 4-7 replicates, except SCA-6 which has only 2 replicates. Another experiment tested the effect of 5 mM dithiothreitol (DTT) on explants during shipping, explants were evaluated by the percent of the petals and staminodes forming calli. The tested cultivars included: SCA-6, CCN-51, INIAPG-038 and Matina 1-6. All of these shipment methods included maintaining the tissues at 4°C and shipping them with or without a 5 mM DTT solution. In total four cultivars were tested using over 1000 staminodes per cultivar across 2-10 replicates per treatment to determine the effects of using DTT.

### Initiation of secondary somatic embryogenesis using primary embryos

To induce secondary somatic embryogenesis, cotyledons were excised from primary somatic embryos and dissected into 4-10 mm segments. These explants were handled according to the method described by Maximova et al. (2002). Secondary somatic embryos were harvested from these cultures while they were on ED3 and used in other processes. Initially, four cultivars, INIAPG-038, CCN-51, SCA-6, and INIAPT-374, were screened for the rates at which primary somatic embryo explants formed secondary somatic embryos. The embryos yielded from these explants were counted at two to three months and each explant ranked into one of four ratings: no embryos produced, 1-4 embryos produced, 5-9 embryos produced, and ≥ 10 embryos produced. In total between 750 and 2700 explants were observed for each cultivar across 5-9 replicates

### Germination and plantlet regeneration

Somatic embryos remained on ED3 medium until they were approximately 15-20 mm in length, at which point they were ready for germination. The germination process began when these somatic embryos were placed onto ED6 medium and were exposed to low levels of light-emitting diode (LED) light, 10-30 umol m-^2^ s-^1^. These embryos were incubated at 27°C and the Petri dishes were then wrapped with 3M Micropore tape (St. Paul, MN) to allow for gas exchange. After two weeks on ED6 the embryos were transferred to EDL media and then maintained on EDL with transfers every two weeks. As the cultures developed, they were slowly exposed to higher LED light intensities, 30-60 umol m-^2^ s-^1^. The germination rates of various cultivars were measured and compared. An embryo was considered germinated if it produced a shoot with a meristem and a root at least 1 cm in length. When the plantlets were 2.5 cm tall, they were transferred to a Phytatray^Tm^ II (Sigma-Aldrich, St. Louis, MO) containing 100 mL of medium and sealed with 3M Micropore tape. The cultivars tested include INIAPT-374, CCN-51 and INIAPG-038. In total between 200 and 900 embryos were germinated per cultivar across 4-9 replicates.

### T-DNA delivery efficiency test

Four cultivars, INIAPG-038, SCA-6, CCN-51, and INIAPT-374 were tested for transient fluorescent protein expression (FPE) after transformation. Two different binary vectors, pDDNPTYFP-1 and pDDNPTYFP-2, were used in cacao transformation experiments (Fig. 1); pDDNPTYFP-1 contains neomycin phosphotransferase II (*nptII*)-GlyLink-yellow fluorescent protein (YFP) translational fusion driven by the cauliflower mosaic virus 35S (CaMV35S) promoter (Fig. 1a) while pDDNPTYFP-2 contains two separate gene cassettes, *nptII* driven by the enhanced CaMV35S promoter and YFP driven by the Nos promoter/TMV Ω enhancer (Fig. 1b). Explants were assessed based on FPE 7 days after *Agrobacterium* infection. Each explant was examined and ranked based on the percentage of its surface area that is expressing YFP. There were six rankings in total: no fluorescent expression, 1-5% FPE coverage, 6-10% FPE coverage, 11-20% FPE coverage, 21-40% FPE coverage, and > 40% FPE coverage. The best treatment was determined by which treatment had the greatest percent of explants in the two highest FPE coverage rankings, however, it was sometimes useful to consider the distribution throughout all the rankings, especially the “no expression” ranking. In total between 450 and 3200 explants were used for each cultivar across 6 to 22 replicates.

**Fig. 1.**
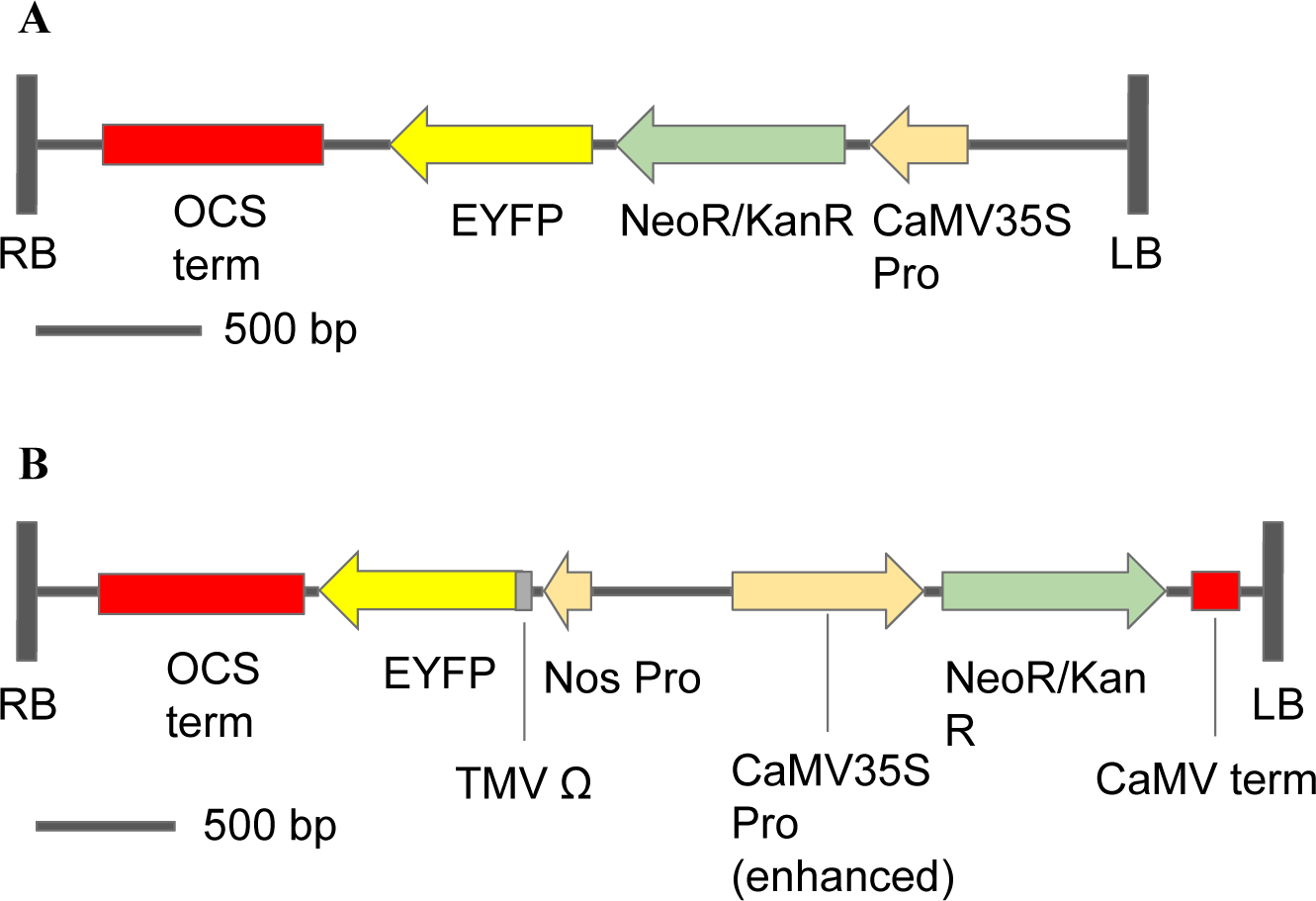
Schematic diagram of two transformation vectors used for cacao transformation. **(a)** pDDNPTYFP-1 is a 3637-bp T-DNA fragment containing *npt*II and *eyfp* translational fusion driven by a CaMV35S promoter. **(b)** pDDNPTYFP-2 is a 4416-bp T-DNA fragment containing a two gene concept where one gene has *eyfp* driven by a Nos promoter/TMV Ω enhancer and the other gene has *npt*II driven by an enhanced CaMV35S promoter

### Transformation and tissue culture improvements in INIAPG-038

Four *Agrobacterium* strains, LBA4404, AGL1, EHA105 and GV3101, containing pDDNPTYFP-1 were tested for T-DNA delivery efficiency. To select the *Agrobacterium* strain, two independent replicates using approximately 400 explants sourced from INIAPG-038 were used. All excised cotyledon tissue from the secondary somatic embryos was collected into a single Petri dish and randomized before being separated and treated with four different *Agrobacterium* strains. Prior to infection, each *Agrobacterium* strain was suspended into the infection medium and then diluted so that each *Agrobacterium* treatment was the same optical density at 600 nm wavelength (OD600). These experiments were evaluated based on FPE coverage at 1 week after infection using the method described previously.

The effect of somatic embryo size on T-DNA delivery was also examined in INIAPG-038. Somatic embryos ranging in size between 4-10 mm were compared to somatic embryos 10-20 mm. In total 700 to 1000 explants for each treatment across 5 to 9 replicates were averaged to determine which embryo sized yielded the greatest T-DNA delivery. Explants were evaluated with the method described previously.

The effect of basal salt concentration, micronutrient concentration as well as the addition of 4.0 uM of CuS0_4_ (5.0 uM final concentration) was examined for their effect on secondary embryo production in INIAPG-038. In total 200 to 400 explants for each treatment across 2 to 5 replicates were averaged to determine which treatments were beneficial compared to a negative control. The explants were evaluated with the method described previously

#### Stable transformation of INIAPG-038

### Agrobacterium infection, co-cultivation, embryogenesis, and germination

Transformations were performed using cotyledon tissue derived from somatic embryos of proper size, color and morphology. The donor material used for the transformation process was secondary somatic embryos, making the transformation a tertiary somatic embryogenic process. To prepare the explants cotyledon tissue from the previously selected secondary embryos was excised then dissected into 4-10 mm^2^ segments. The infection step was performed by suspending the explants in 10-20 mL of liquid medium harboring *Agrobacterium* contained within a sealed 50 mL test tube. The infection medium consisted of MS basal salt and vitamins containing 200 uM acetosyringone, 1 mg/L indole acetic acid and 2 mg/L zeatin. The *Agrobacterium* was thoroughly suspended in the liquid infection medium and the OD600 was measured prior to use and diluted to an OD600 of 0.400. After 1-2 hours of agitation, the *Agrobacterium* infected cotyledon explants were transferred to a SCG based co-cultivation medium and incubated at 21°C for three to four days before the *Agrobacterium* was washed away. The cotyledon tissue was then transferred to a modified SCG medium containing 200 mg/L cefotaxime and 200 mg/L timentin. The tissue remained on this antibiotic containing SCG for between two and four weeks before transfer to ED4 medium containing the same antibiotics. Antibiotics remained in the medium whenever there were explants that had been directly exposed to *Agrobacterium*. However, later in the process during the germination of quaternary somatic embryos the antibiotics were removed from the medium. The fourth transformation included the use of 100 mg/L of kanamycin for chemical selection. After two weeks of resting on SCG the explants were moved to ED4 containing both antibiotics and kanamycin. This kanamycin concentration was maintained for 1 month then increased to 150 mg/L while the explants were on ED3.

The *Agrobacterium*-infected tissue underwent indirect somatic embryogenesis and was observed using a fluorescent microscope multiple times between one and four months during the formation of embryos. After the identification of a transgenic event through the use of the fluorescent microscope the transgenic, tertiary somatic embryos (TSEs) were removed from the remainder of the non-transgenic tissue and isolated on ED3. To generate multiple quaternary somatic embryos (QSEs) an additional secondary somatic embryogenesis step was used. QSE germination and plantlet regeneration were conducted as described above.

### PCR analysis

A CTAB DNA extraction protocol, described by Murray and Thompson (1980) was used to extract genomic DNA from leaf materials of nontransgenic INIAPG-038 and transgenic INIAPG-038 events. To test the presence of *npt*II in transgenic plants, the primer set, NPTII 3F (5’-CAAGATGGATTGCACGCAGGTT-3’) and NPTII 4R (5’-TAGAAGGCGATGCGCTGCGAAT-3’) was used while the presence of *eyfp* was determined using the primer set, EYFP 3F (5’-TAAACGGCCACAAGTTCAGCG-3’) and EYFP 4R (5’-AGGACCATGTGATCGCGCTTC-3’). For each PCR reaction mixture, the following reagents were used: 25 uL DreamTaq PCR Master Mix (2X) (Thermo Fisher Scientific, Grand Island, NY), 1.0 uL forward primer at 10 uM, 1.0 uL reverse primer at 10 uM, 21 uL H_2_0, and 2.0 uL genomic DNA at ∼100 ng/uL, for a total volume of 50 uL. The PCR was carried out with the following thermal cycler programming: hold at 95°C for 3 min, 16 cycles of 94°C for 30 sec, 58°C for 30 sec (−0.5°C/cycle), 68°C for 1 min, 50 cycles of 94°C for 30 sec, 54°C for 30 sec, 69°C for 1 min; and then elongation at 68°C for 7 min, and remained at 4°C for infinite. For each PCR reaction, 20 uL were loaded onto a 0.8% agarose gel for electrophoresis.

### Fluorescent visualization

Fluorescent images of embryonic and plantlets of transgenic INIAPG-038 events were visualized with a fluorescent Leica M165 FC stereomicroscope, equipped with Leica DFC7000 T (JH Technologies, Fremont, CA); using two microscopic filters, brightfield and ET YFP with 514 nm excitation and 527 nm emission. The microscope is linked to a camera imaging software, Leica Application Suite version 4.9, which was used to capture the fluorescent images. Screening of fluorescent activity was measured at 30x magnification.

### Transplantation to soil and acclimatization

After embryo germination while the plantlets are 10 to 13 cm in height and have six to ten leaves they were transferred to soil. The plantlets were transferred into a well-draining soil, mixing equal parts sunshine mix and perlite was found to work well. When transplanting the agar was removed by hand and by dipping the roots into water. The plantlets were then planted into the pre-moistened soil mix and immediately covered. Afterwards, the plantlets were placed into a tray and covered with a humidity dome with all the vents closed. On top of the humidity dome shades were placed to reduce the light intensity to 60 umol m-^2^ s-^1^. Over the course of a week the vents on the humidity dome were opened and the shades were removed. After this acclimatization process the plants were able to survive in the growth chamber with an ambient environment of 27°C, 60-70% RH and 120-180 umol m-^2^ s-^1^ light intensity.

### Grafting of transgenic scions onto in vitro germinated rootstock

Wedge-shaped grafting was done as previously described by Bezdicek (1982) with modifications. Cacao pods were surface sterilized using 10% bleach solution for 15 minutes, then the pod was cut open and the seeds removed. The thick pulp was then removed from the seeds. The seeds were further surface sterilized using 10% bleach solution for 15 minutes. The sterilized seeds were then soaked in sterile water for 72 hrs. Afterwards, these pre-germinated seeds were planted into autoclave-sterilized peat pellets (Jiffy Products of America Inc. Lorain, OH, USA) soaked in a liquid EDL medium. The seedlings were kept in Phytatrays and cultured similar to tissue culture-derived plantlets.

When both a transgenic scion with poor roots and a wild-type *in vitro* root stock with strong, healthy roots were available, grafting was performed using a “V” shape cut similar to a saddle or cleft graft on the seedling and two diagonal cuts exposing the cambium layer of the transgenic scion. The scion and rootstock were then combined and secured using both a paper clip and a 1 cm segment of a polypropylene straw cut down the side to resemble a “C”. All the cuts were performed and secured using sterilized tools and materials. The grafts were then sealed in Phytatrays with micropore tape and liquid EDL medium was periodically added to maintain proper moisture. After the scion and rootstock fused, they were transplanted to the soil using the previously outlined method.

## Results and Discussion

To establish a successful, highly efficient transformation protocol, a tissue culture system spanning the initiation of floral material to the regeneration of plantlets into soil has been tested and improved in cacao. Multiple cultivars were screened for their response to (1) primary embryo induction from floral material, (2) secondary embryo production from cotyledon tissue derived from primary somatic embryos, (3) embryo germination/regeneration to plantlets, and (4) transfer DNA (T-DNA) delivery efficiency via *Agrobacterium*. These four factors were used to determine how amenable various cultivars would be for genetic transformation.

### Tissue culture response and INIAPG-038 improvements

Petals and staminodes were used to initially test somatic embryogenesis in 14 cacao cultivars. Staminode explants generally formed embryos at a higher frequency than petal explants in all cultivars except SCA-6 where they were equal. (Fig. 2). This is consistent with previous observations made by Tan et al. (2003) and Traore et al. (2006). However, Garcia et al. (2016) observed that petals produced on average more embryos per explant. The cultivar with the highest percentage of floral explants forming embryos was SCA-6 with 44%/44% of petal/staminodes forming embryos (Fig. 2). The next best performing cultivars were CCN-51, INIAPG-038, and INIAPT-374 with petal/staminodes producing embryos at a rate of 6%/29%, 7%/15%, and 4%/21%, respectively. The remainder of the cultivars had less than 10% of floral explants producing embryos. INIAPG-069 was most recalcitrant and Matina 1-6, GS 36, INIAPT-484, and INIAPG-029 were slightly less recalcitrant.

**Fig. 2.**
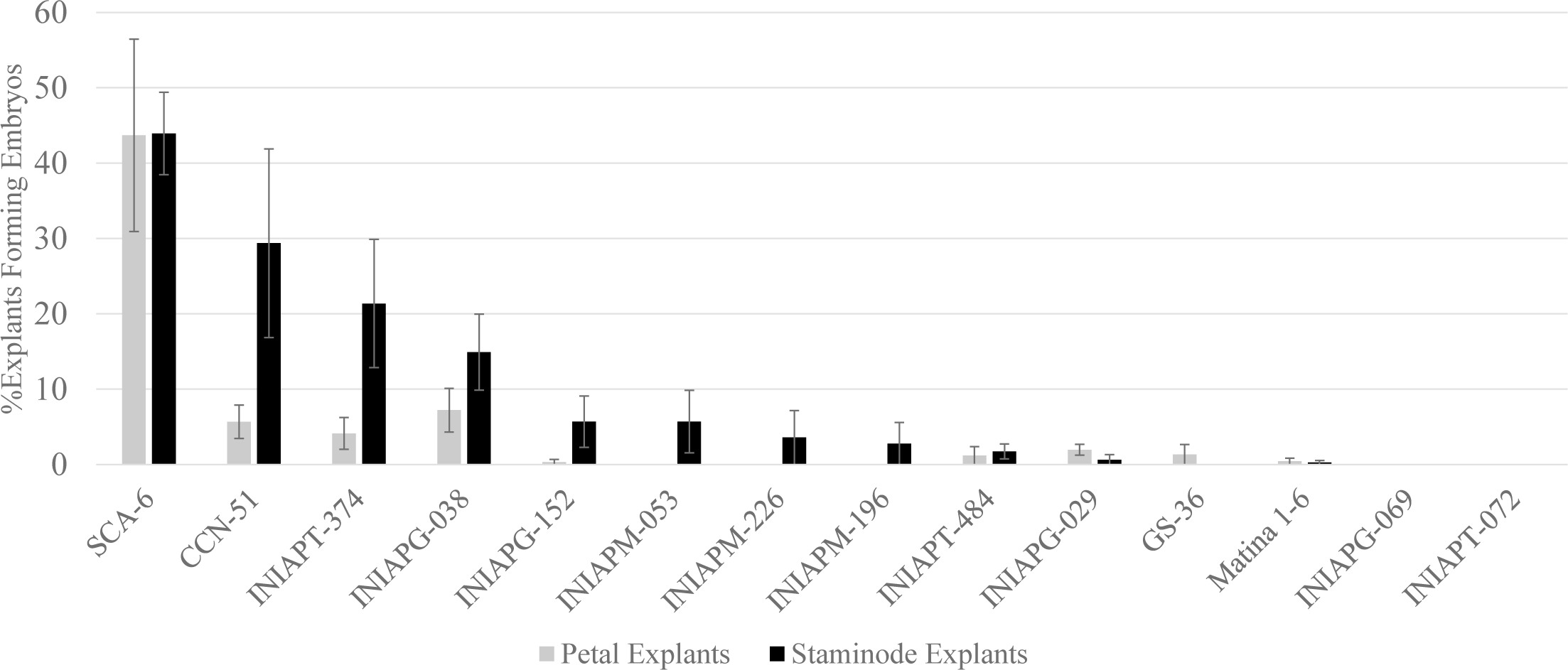
Cumulative averages of somatic embryo formation from flower materials for 14 different cacao cultivars. Petal and staminode explants were used to induce primary somatic embryos. Each histogram represents the mean level (±standard error) from between 2 and 7 replicates

There was a significant oxidation, browning, and viability loss of donor flower materials even during overnight shipping from USDA ARS Subtropical Horticultural Research Station (SHRS) in Miami, FL, to IGI, Berkeley, CA, USA (Fig. 3a). Tissue browning is a typical phenomenon observed in cultures obtained from mature explants of the woody plants, and it affects callus growth and adventitious shoot regeneration (Dutta Gupta, 2010). It was previously reported that an antioxidant DTT improved callus formation, shoot regeneration, and rooting by inhibiting the accumulation of peroxidase in Virginia pine (*Pinus virginiana* Mill.) (Wei et al. 2004). Therefore, DTT was tested to prevent plant tissue oxidation and necrosis during the shipment. The shipment of flowers overnight in a solution of 5 mM DTT was found to dramatically improve the viability of the immature flowers with less oxidation and browning of flowers, compared to no DTT treatment (Fig. 3). In the cultivar CCN-51 an increase of 740% more staminodes formed callus compared to the no DTT treatment (Fig. 4). The three other cultivars, CCN-51, INIAPG-038 and SCA-6, tested showed a similar increase in the formation of callus with the 5 mM DTT shipping treatment.

**Fig. 3.**
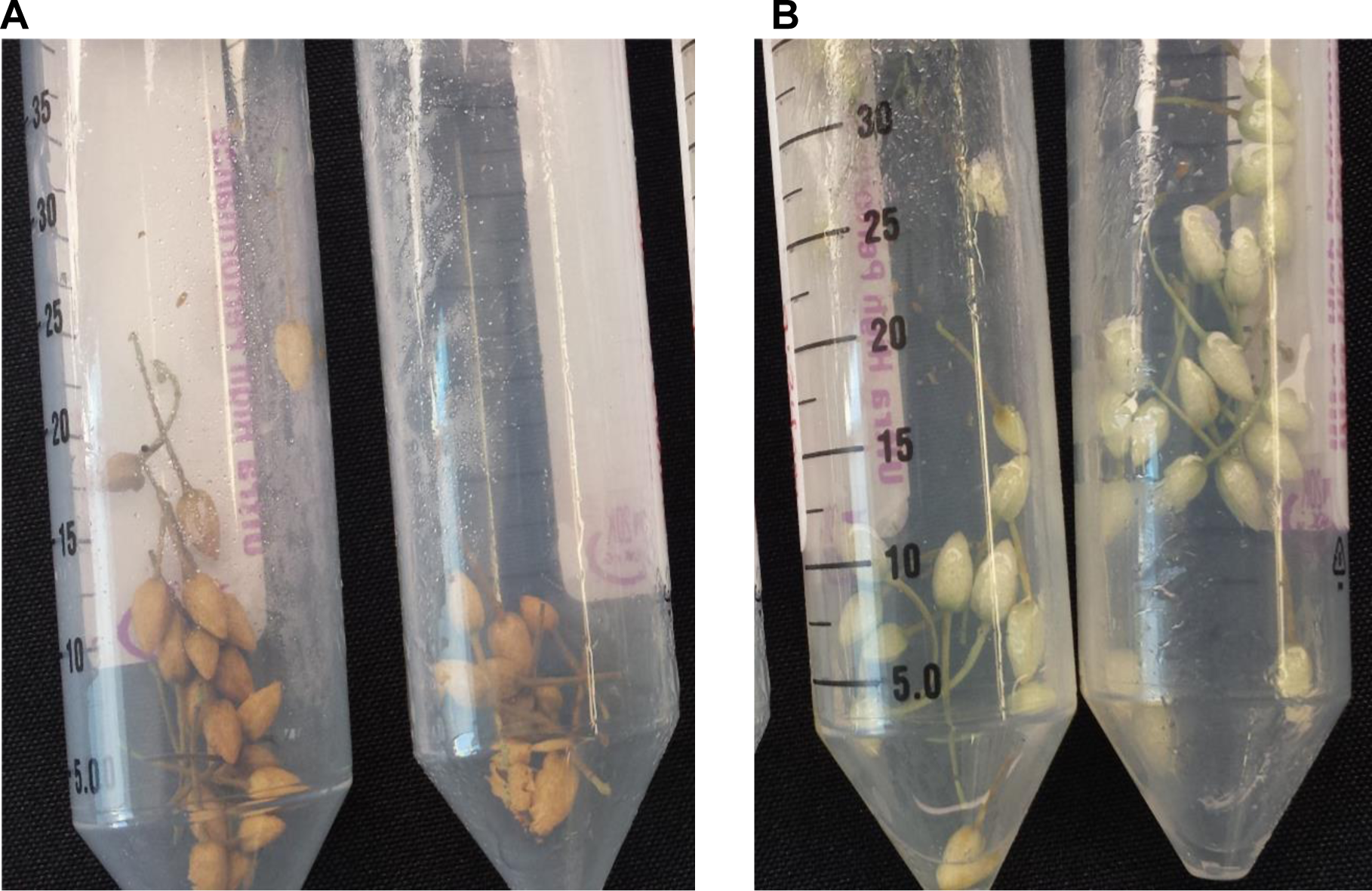
INIAPG-038 flowers shipped dry without DTT **(a)** and in 5 mM DTT **(b)**. Both treatments were contained in 50 mL conical tubes and kept at 4°C overnight

**Fig. 4.**
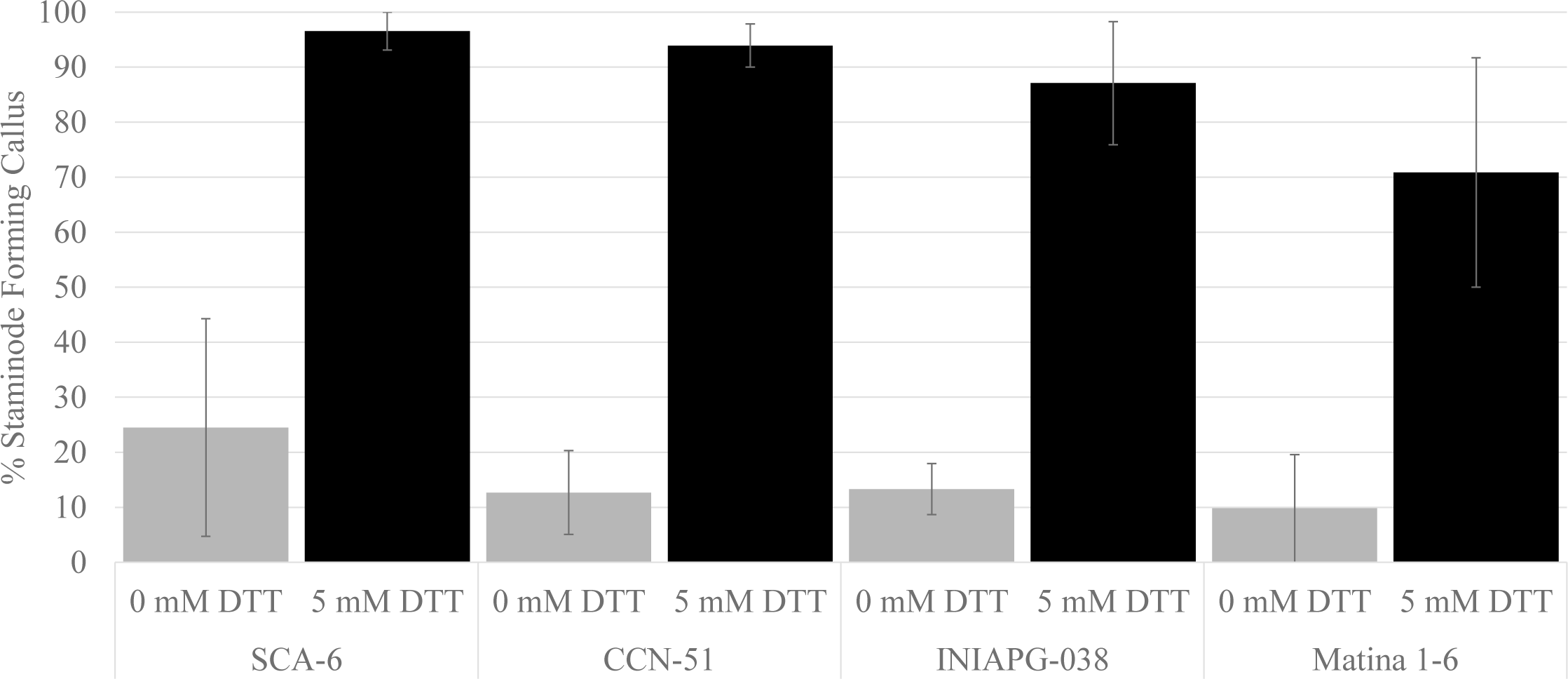
Effect of 5 mM DTT solution on flower material tissue culture response. Immature flowers were either shipped in a 5 mM DTT Solution or dry without a DTT solution. Both treatments were contained in 50 mL conical tubes and kept at 4°C overnight. Each histogram represents the mean level (±standard error) from between 2 and 10 replicates

The four cultivars, SCA-6, CCN-51, INIAPT-374 and INIAPG-038, showing the best primary somatic embryo formation (Fig. 2) were further tested for their secondary somatic embryo production (Fig. 5). It was observed that INIAPG-038 had the highest percentage of explants producing greater than 10 embryos at 6.2%. It also had the greatest number of explants producing between 5 and 9 embryos at 12.6%. The next highest performing cultivars were SCA-6, CCN-51 and INIAPT-374 with 4.5%/10.2%, 3.9%/9.7% and 1.0%/2.8% of explants forming greater than 10 embryos/explants forming between 5 and 9 embryos, respectively. In total three cultivars were tested for the rate at which embryos would regenerate into plantlets (Fig. 6). The highest performing cultivar was INIAPT-374 with embryo regeneration rates of 33%. CCN-51 and INIAPG-038 embryos regenerated to plantlets at a rate of 28% and 22%, respectively.

**Fig. 5.**
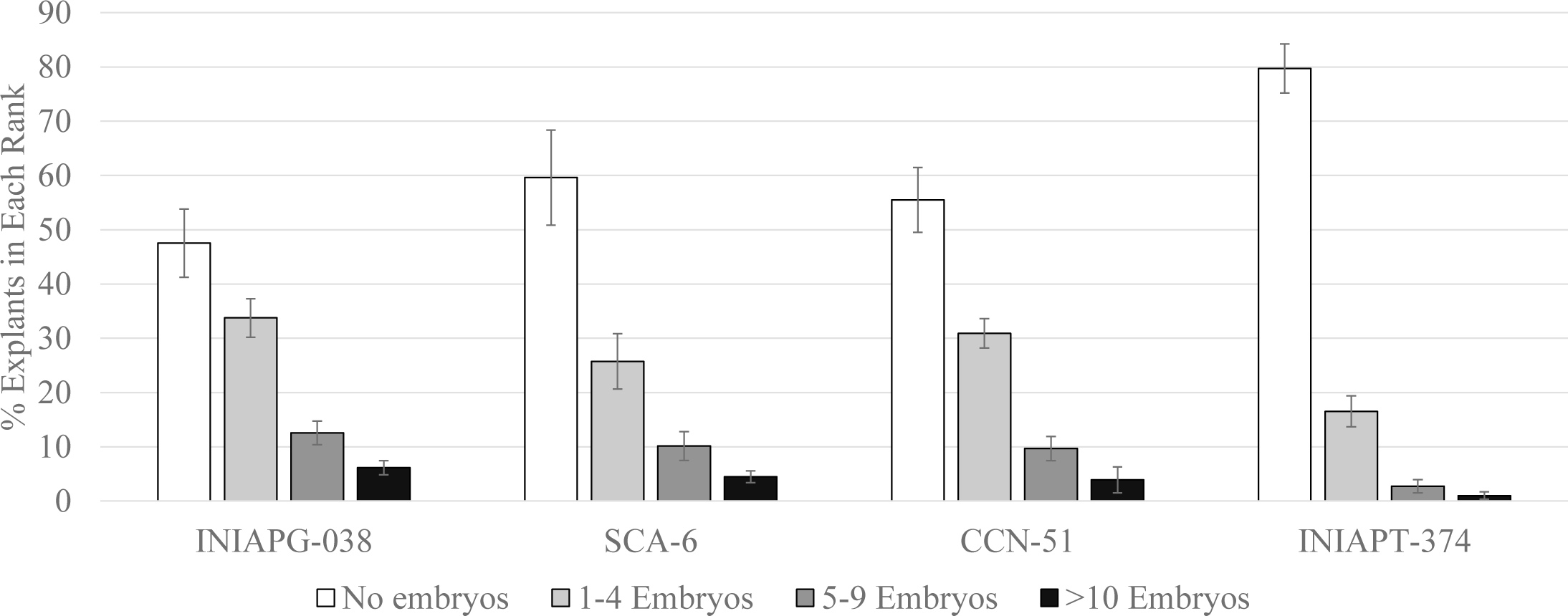
Cumulative averages of secondary somatic embryogenesis for selected cultivars. Cotyledon explants were observed in four ranks based on the number of embryos they produced: No embryos produced, 1 to 4 embryos produced, 5 to 9 embryos produced, and greater than 10 embryos produced. Each histogram represents the mean level (±standard error) from between 5 and 9 replicates

**Fig. 6.**
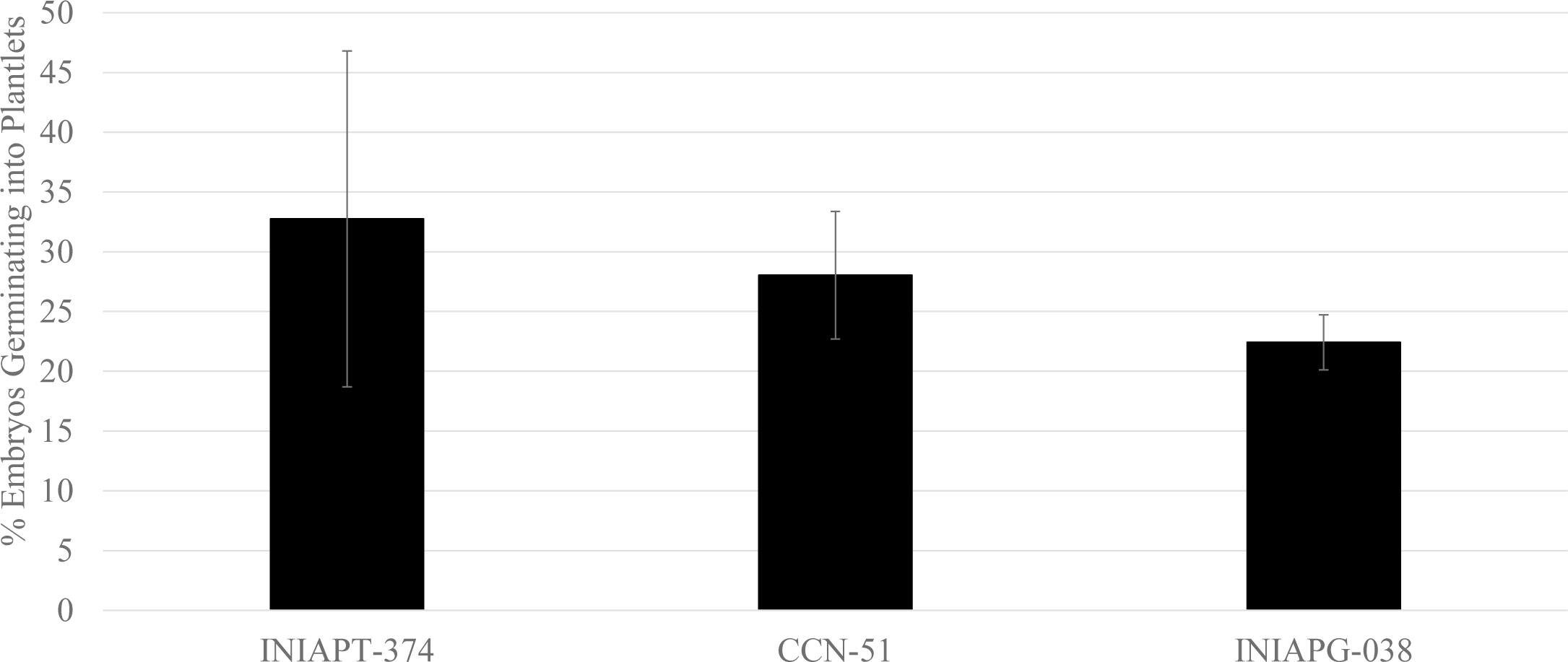
Cumulative averages of plantlet regeneration for selected cultivars. In total 256 INIAPT-374, 227 CCN-51 and 868 INIAPG-038 embryos were used to test regeneration rate. Each histogram represents the mean level (±standard error) from between 4 and 9 replicates. Mature embryos were exposed to increasing levels of light while on ED6 then EDL media and counted as germinated if they produced a 1 cm shoot with meristem and a root of 1 cm

With regard to experiments conducted to improve secondary somatic embryogenesis in INIAPG-038, it was observed that using a 2-fold concentration of woody plant medium (WPM) basal salts (Lloyd and McCown 1981) plus 1-fold vitamins significantly increased embryo production (Fig. 7). The 1-fold WPM basal salts plus vitamins negative control had 2.5% of explants forming greater than 10 embryos and 12.0% forming between 5 and 9 embryos, whereas 2-fold basal medium resulted in 12.3% and 16.5%, respectively. SCG medium, WPM-based medium, contains 1.0 uM CuSO4, 10-fold higher than MS medium (Murashige and Skoog 1962). A five-fold copper level (5.0 uM) in SCG medium also resulted in 10.9% of explants producing greater than 10 embryos and 23.4% of explants forming 5 to 9 embryos in INIAPG-038 (Fig. 7). This is consistent with the previous conclusion that the callus quality, callus growth and regenerability can be improved by increasing level of the micronutrient copper in barley and oat (Dahleen 1995; Cho et al. 1998, 1999).

**Fig. 7.**
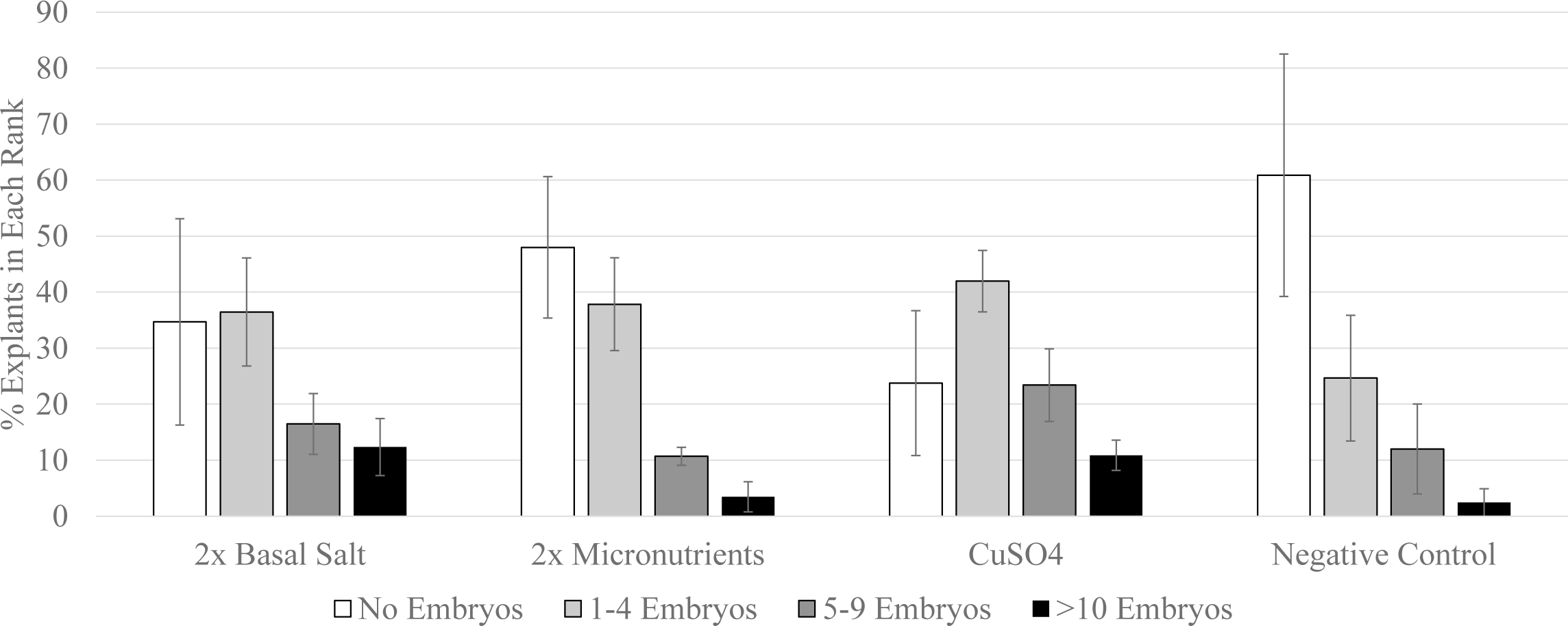
Effect of two-fold WPM basal salt, two-fold WPM micronutrients, and addition of 4.0 uM cupric sulfate on secondary somatic embryogenesis in INIAPG-038. Cotyledon explants were observed in four ranks based on the number of embryos they produced: No embryos produced, 1 to 4 embryos produced, 5 to 9 embryos produced, and greater than 10 embryos produced. Each histogram represents the mean level (±standard error) from between 2 and 5 replicates

### T-DNA Delivery Efficiency Improvements and Optimizations

Experiments for the T-DNA delivery efficiency test were conducted using pDDNPTYFP-1 containing *npt*II::*eyfp* in the 4 cultivars with good tissue culture response: INIAPG-038, CCN-51, SCA-6 and INIAPT-374. Out of them, INIAPG-038 had the highest T-DNA delivery efficiency (Fig. 8). Whereas CCN-51 had the highest number of explants ranked as >40% and 20-40% coverage at 7.1% and 7.2% respectively, INIAPG-038 was comparable at 6.3% and 5.7%. INIAPG-038 however only had 34.7% of explants with no FPE while CCN-51 was at 39.5%.

**Fig. 8.**
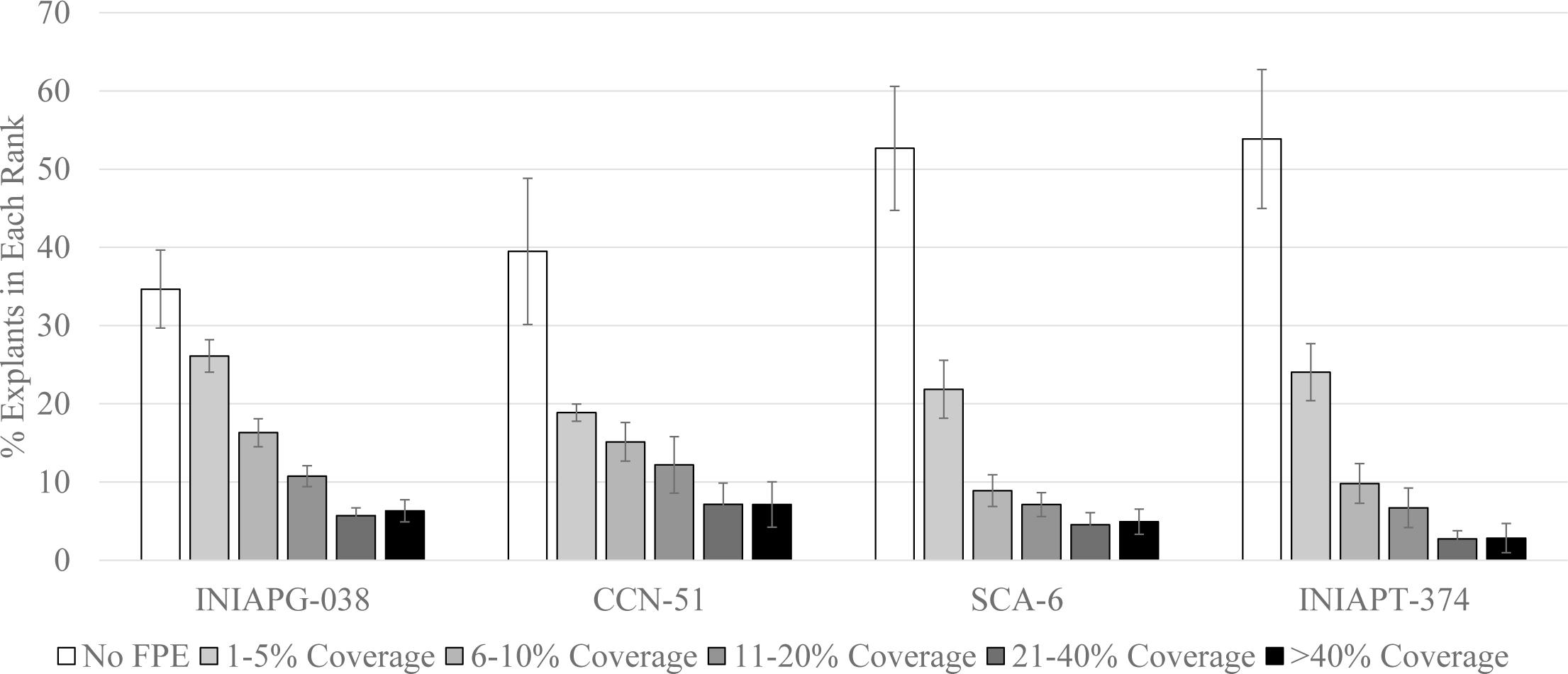
Cumulative averages of transient YFP expression for selected cacao cultivars. Cotyledon explants were observed in six ranks based on FPE coverage: No Expression, 1-5% FPE coverage, 6-10% FPE coverage, 11-20% FPE coverage, 21-40% FPE coverage and greater than 40% FPE coverage. Each histogram represents the mean level (±standard error) from between 6 and 22 replicates

Of the four *Agrobacterium* strains tested, AGL1 produced the highest transient FPE coverage, compared to 3 other *Agrobacterium* strains (Fig. 9). EHA105 had the 2nd highest transient FPE coverage, but LBA4404 and GV3101 performed poorly and produced no explants with >40% transient FPE coverage. This was seen previously in stable transformation of an elite maize inbred (Cho et al. 2014). However, other plant species or explants favored different *Agrobacterium* strains such as LBA4404 used in tobacco transformation (Bakhsh et al. 2014) and GV3101 used in tomato transformation (Chetty et al. 2013).

**Fig. 9.**
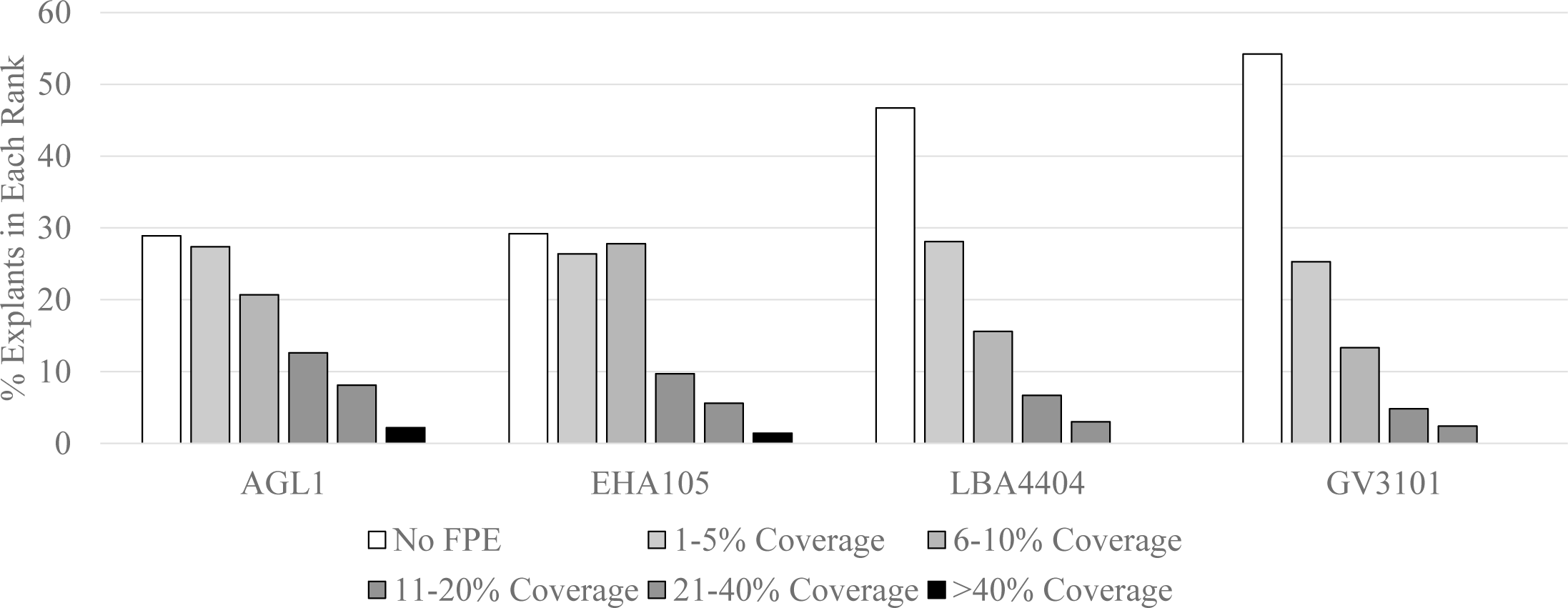
Effect of *Agrobacterium* strain on transient YFP expression in INIAPG-038. Cotyledon explants were observed in six ranks based on FPE coverage: No Expression, 1-5% FPE coverage, 6-10% FPE coverage, 11-20% FPE coverage, 21-40% FPE coverage and greater than 40% FPE coverage. Each histogram represents the mean level

With regard to the INIAPG-038 explant type used for cacao transformation, we observed that cotyledon tissue derived from primary or secondary somatic embryos had the highest transient FPE, while hypocotyl and radicle explants demonstrated very little transient FPE and embryo production after *Agrobacterium* infection (data not shown). Previously cotyledon tissues derived from mature embryos of PSU SCA-6 were used for transformation target (Maximova et al. 2003, 2005; Florez et al. 2015; Shires et al 2017). It was observed that the embryo size of INIAPG-038 also affected transient FPE after transformation (Fig. 10). Cotyledon tissues derived from immature somatic embryos ranging in size between 4-10 mm outperformed those derived from mature embryos ranging in size between 10-20 mm. It was observed that embryos ranging in the 4-10 mm range produced explants with >40% FPE coverage at a rate of 12.3% and explants with 21-40% FPE coverage at 9.1% whereas 10-20 mm embryos only produced explants in those ranks at 1.9% and 4.0%, respectively.

**Fig. 10.**
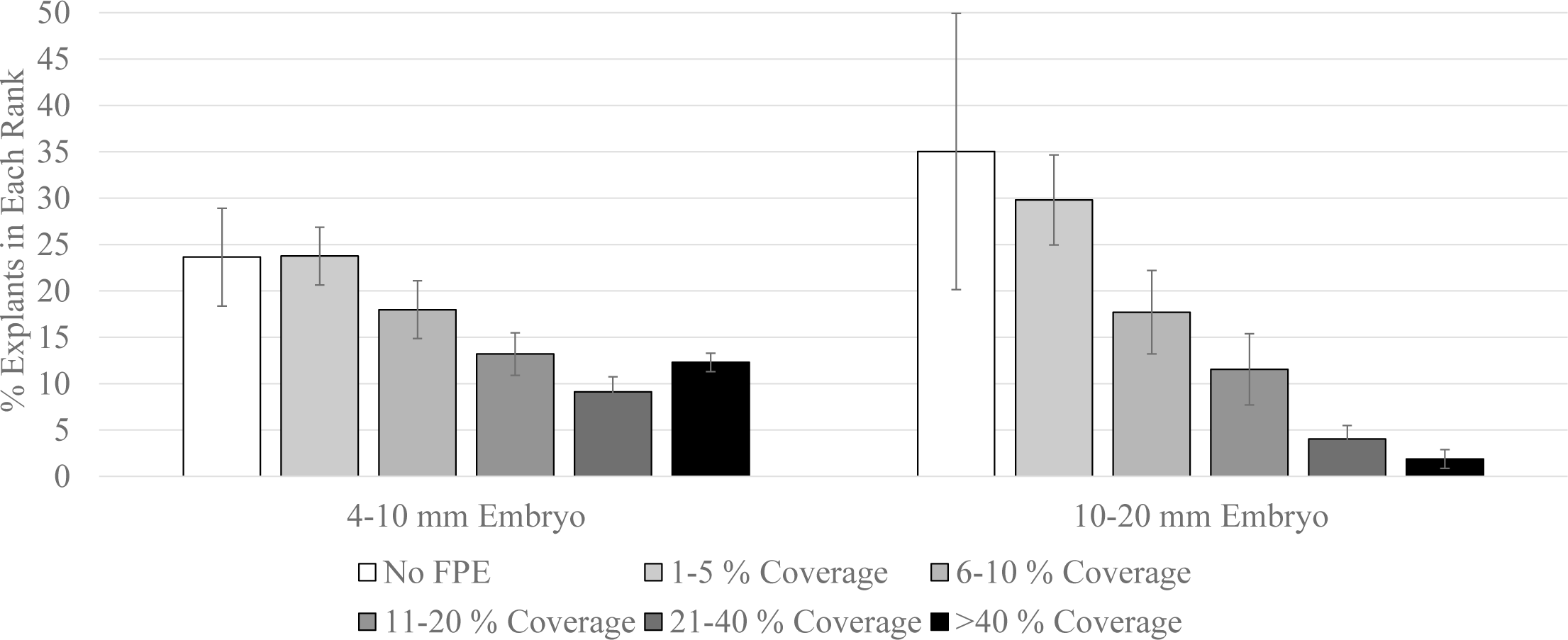
Effect of embryo size on transient YFP expression in INIAPG-038. Cotyledon tissue derived from somatic embryos was used for experiments. Cotyledon explants were observed in six ranks based on FPE coverage: No FPE, 1-5% FPE coverage, 6-10% FPE coverage, 11-20% FPE coverage, 21-40% FPE coverage and greater than 40% FPE coverage. Each histogram represents the mean level (±standard error) from between 5 and 9 replicates

### Stable Transformation of INIAPG-038

INIAPG-038 was selected for further improvements to tissue culture response, T-DNA delivery efficiency, and eventually for stable transformation because it performed the best in both secondary somatic embryogenesis and T-DNA delivery. Although out of the 4 best cultivars from primary somatic embryogenesis screening it performed less efficiently in both primary somatic embryogenesis and germination/regeneration (Figs. 2 and 6), these two steps are not as crucial for transformation and are not seen as potential bottlenecks. In addition, INIAPG-038 is a highly productive cultivar with tolerance to frosty pod disease and witches broom disease which weighed on the decision to select it. Although not selected for transformation, CCN-51 shows promise for transformation because it performed well in terms of both tissue culture response and T-DNA delivery. CCN-51 is another highly productive cultivar that is widely planted in Latin America. Previous transformations were conducted on PSU SCA-6 (Maximova et al. 2003, 2006; Florez et al. 2015; Shires et al. 2017) because it is an amenable cultivar with high somatic embryogenesis and genetic transformation potential (Maximova et al. 2006). This Forastero cultivar also demonstrates high production of primary somatic embryos when compared to Trinitario cultivars (Traore et al. 2006). Our decision to focus on INIAPG-038 was heavily weighed on the importance of the cultivar as well as the somatic embryogenic potential (Figs. 2 and 5) and the T-DNA delivery efficiency (Fig. 8). Stable transformation experiments in INIAPG-038 were conducted by infecting cotyledon tissues derived from the secondary somatic embryos with AGL-1 containing pDDNPTYFP-1 or pDDNPTYFP-2 (Fig. 1). Transient YFP expression was clearly observed 5-7 days after *Agrobacterium* infection (Fig. 11A). Four separate successful stable transformation experiments in INIAPG-038 were conducted by YFP visual marker selection. Kanamycin selection was not tight enough and nontransgenic embryo tissue still formed on the cotyledon explants even with a high level of kanamycin. In the method described by Maximova et al. (2003) 50 mg/L of geneticin (G418) for *npt*II was used which greatly reduces the formation of non-transgenic tissues. Eight independent YFP-expressing somatic embryos of INIAPG-038 (Fig. 11B) were generated using cotyledon tissues derived from 82 somatic embryos; transformation frequency at the T_0_ tissue level was 9.8% (8/82) (Table 1). Of these eight independent transgenic events, three had decent embryo morphology. The transgenic embryos produced through our experiments have been few in number and the average rate of embryo germination for INIAPG-038 remained 22.4% (Fig. 6). So, to increase the likelihood of regenerating these events the cotyledon tissue of the transgenic, an additional embryogenic cycle was conducted to generate multiple QSEs before germination. All three independent transgenic TSEs were capable of producing multiple QSEs (Figs. 11C and D) and were capable of regeneration into plantlets (Fig. 11E), resulting in a transformation frequency of 3.7% (3/82) (Table 1). Similarly, an additional embryogenic step after the production of transgenic cacao embryos was also used by Maximova et al. (2003). The recalcitrant nature of cacao and the low rate at which embryos convert into plants, below 30%, makes direct regeneration of transgenic embryos a risky endeavor, most likely leading to the loss of the majority of transgenic events. Using the quaternary somatic embryogenesis process a single transgenic embryo could be proliferated into multiple somatic embryos. The hypocotyl and plumule were preserved throughout this process and maintained on ED3 medium. Additional embryogenic cotyledon tissue developed from the transgenic hypocotyl so the quaternary somatic embryogenesis process could be conducted multiple times. However, once the original transgenic tissue matured, it was transferred to ED6 for germination. With the quaternary somatic embryogenesis all three of the transgenic events, #1, 2 and 3, proliferated into 68, 16, and 95 total embryos, respectively (Table 1). One (event #3) of the three transgenic embryos was chimeric and nontransgenic QSEs were discarded based on YFP expression. Then 15, 4, and 20 of those embryos germinated into plantlets, respectively (Table 1), giving an average of 21.8% rate of transgenic embryo conversion to plantlets. Eleven, 3, and 3 plants of each event were successfully acclimatized in soil, respectively, so far (Table 1and Fig. 11F).

**Table 1.**
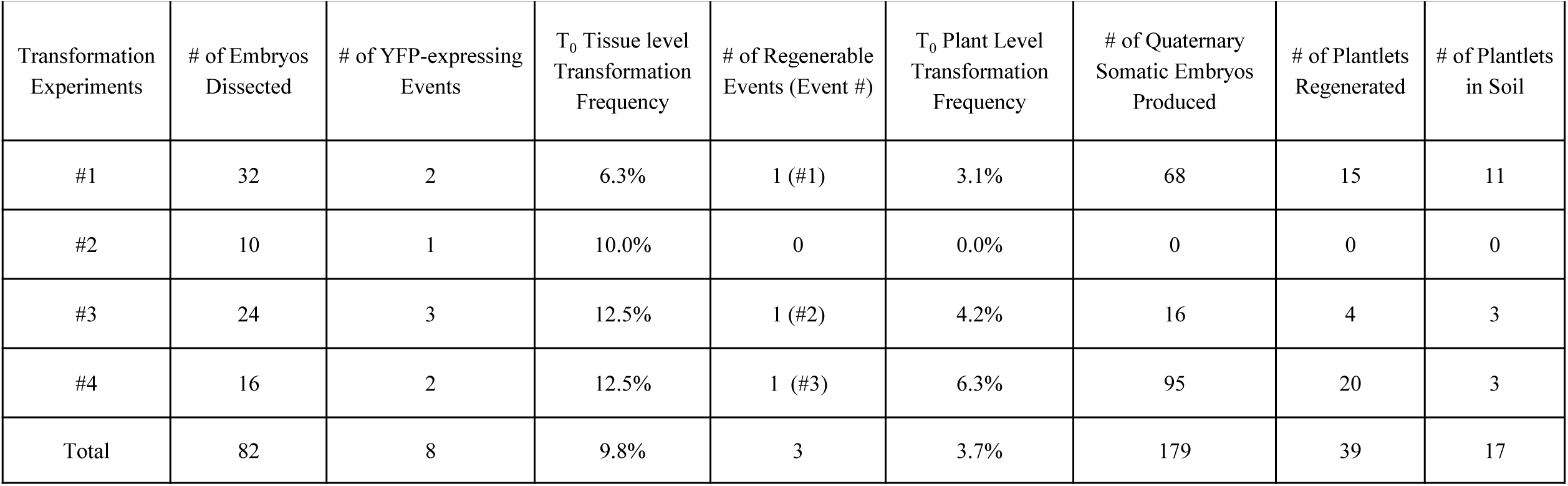
Stable transformation frequencies of INIAPG-038

| Transformation Experiments | # of Embryos Dissected | # of YFP-expressing Events | T<sub>0</sub> Tissue level Transformation Frequency | # of Regenerable Events (Event #) | T<sub>0</sub> Plant Level Transformation Frequency | # of Quaternary Somatic Embryos Produced | # of Plantlets Regenerated | # of Plantlets in Soil |
| --- | --- | --- | --- | --- | --- | --- | --- | --- |
| #1 | 32 | 2 | 6.3% | 1 (#1) | 3.1% | 68 | 15 | 11 |
| #2 | 10 | 1 | 10.0% | 0 | 0.0% | 0 | 0 | 0 |
| #3 | 24 | 3 | 12.5% | 1 (#2) | 4.2% | 16 | 4 | 3 |
| #4 | 16 | 2 | 12.5% | 1 (#3) | 6.3% | 95 | 20 | 3 |
| Total | 82 | 8 | 9.8% | 3 | 3.7% | 179 | 39 | 17 |

**Fig. 11.**
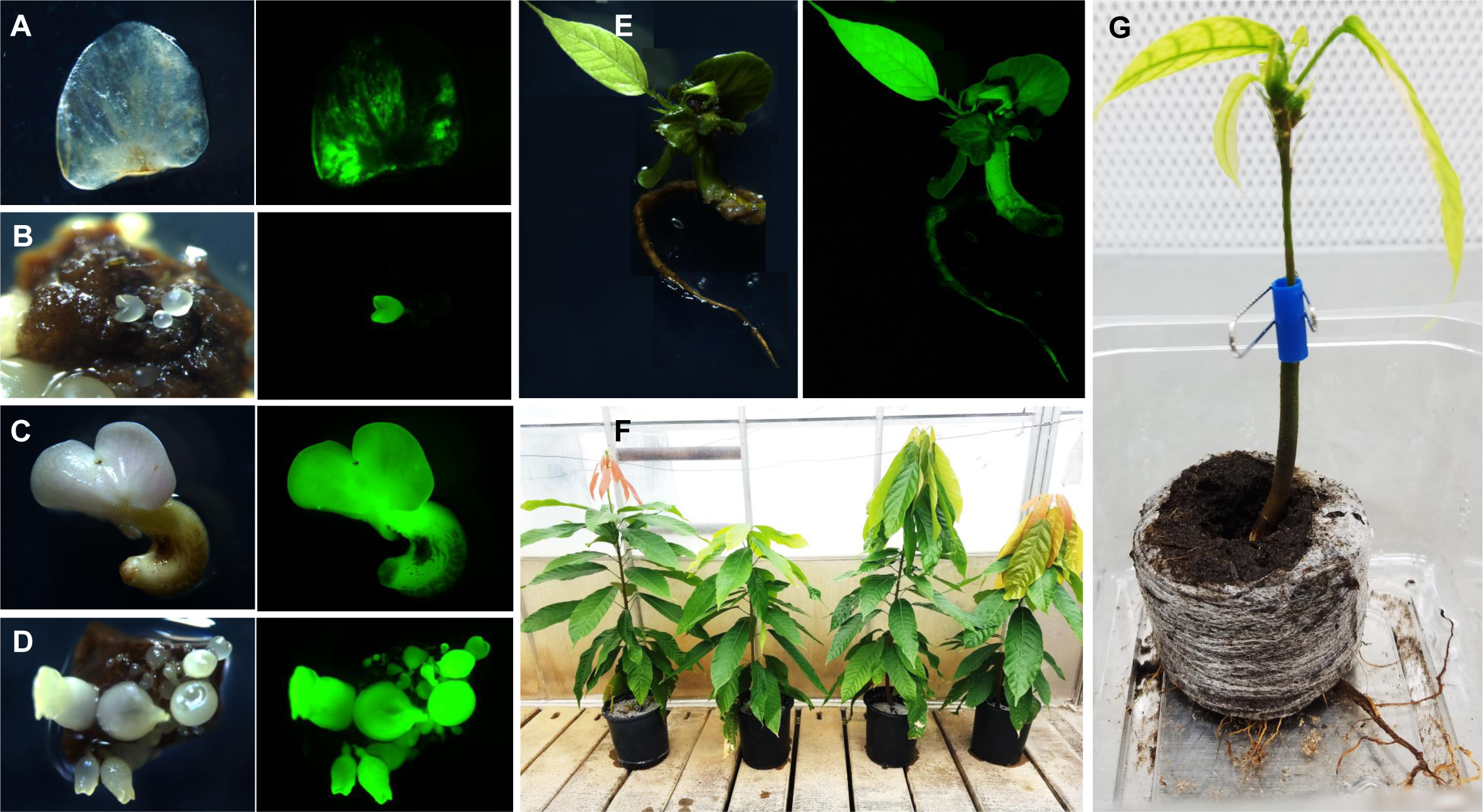
Transient YFP expression and stable transgenic event development in INIAPG-038. **(a)** Cotyledon expressing transient YFP. **(b)** YFP-expressing globular embryo. **(c)** YFP-expressing germinating embryo. **(d)** YFP-expressing quaternary somatic embryos. (e) YFP expressing plantlet. **(f)** A nontransgenic (leftmost) and 3 transgenic INIAPG-038 plants in the greenhouse. **(g)** A grafted transgenic INIAPG-038 plantlet in a Jiffy peat pellet; a transgenic INIAPG-038 scion with poor roots was grafted onto a wild-type rootstock

Some regenerated transgenic INIAPG-038 plantlets had slow or poor development of roots and these could not survive in soil, so grafting was attempted to resolve this issue. Plantlets with poorly developed root systems routinely die during transplanting to soil, there is also a risk of damaging well-established root systems during agar removal. However, by grafting transgenic scions *in vitro* prior avoids these two issues since a robust well-rooted wild-type seedling is used and the entire intact root system within a sterilized Jiffy peat cube can be placed into the soil without disturbing the roots (Fig. 11G). In total, six transgenic scions with poor roots were grafted onto rootstock, four of these survived. In addition, grafting significantly facilitated the use of transgenic shoots that may have otherwise not formed roots. To our knowledge, this is the first report of this technique in transgenic cacao. The use of grafting to overcome poor *in vitro* rooting and improve transformant recovery was also used previously in safflower and cotton (Jin et al. 2006; Belide et al. 2011). Our recent results showed that sonication treatment significantly increased T-DNA delivery efficiency. Twelve seconds of sonication increased the percent of explants exhibiting >21% FPE by 416% (Fig. S1), suggesting that there is a lot of potential for improvement to stable transformation in INIAPG-038.

The presence of transgenes, *eyfp* and *npt*II, was confirmed by PCR analysis. The amplified product of 595 bp corresponding to the internal fragment of *eyfp* gene was observed from genomic DNA of all nine tested transgenic plants derived from three independent events using *eyfp* gene-specific primers, EYFP 3F and EYFP 4R (Fig. 12A). An amplified fragment of 761 bp was also observed from all tested transgenic plants derived from three independent events using *npt*II gene-specific primers, NPTII 3F and NPTII 4R (Fig. 12B). No PCR-amplified *eyfp* and *npt*II fragments were detected in the wild-type control plant (Figs. 12A and B) as expected.

**Fig. 12.**
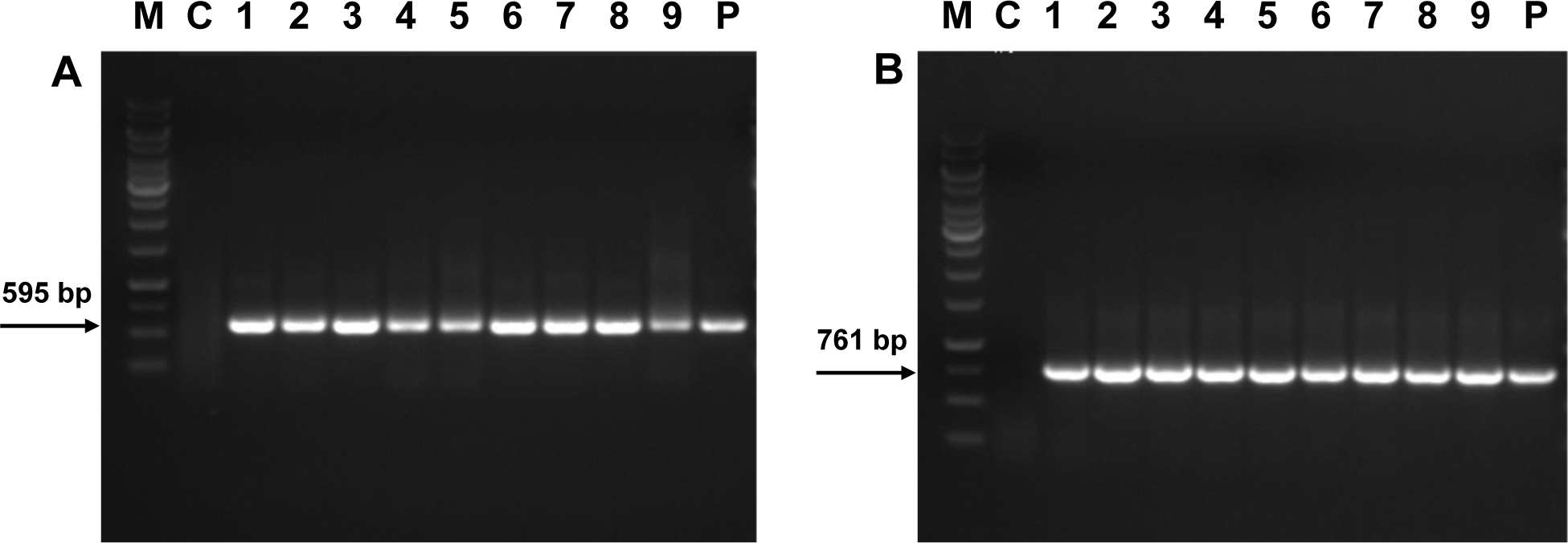
PCR analysis of genomic DNA from three different transgenic INIAPG-038 events. Genomic DNA was extracted from leaf tissues of one nontransgenic control and three plants of each independent transgenic event. *M* GeneRuler 1-kb DNA ladder, *C* Nontransgenic INIAPG-038 control, *Lanes 1-3* Transgenic INIAPG-038 Event #1-1, #1-2 and #1-3, *Lanes 4-6* Transgenic INIAPG-038 Event #2-1, #2-2 and #2-3, *Lanes 7-9* Transgenic INIAPG-038 Event #3-1, #3-2 and #3-3, *P* pDDNPTYFP-2 as a positive control. **(a)** Amplified 595-bp fragment products using *eyfp* gene-specific primers, EYFP 3F and EYFP 4R. **(b)** Amplified 761-bp fragment products using *npt*II gene-specific primers, NPTII 3F and NPTII 4R

In conclusion, we have established a successful *Agrobacterium*-mediated transformation system for the production of transgenic cacao plants using INIAPG-038, a high-yielding, disease-resistant cultivar. Previously only PSU SCA-6 has been reported for stable transformation. The results in this study can be applied to improve tissue culture response and transformation frequency in other cacao cultivars.

## Supporting information

Jones et al_Supplemental_Fig1.pdf

## Acknowledgements

We are grateful to Alexandra Tempeleu and Brian Margolesky (Mars Cacao Laboratory, Miami, FL, USA) for kindly shipping cacao flowers. This work was supported by Mars, Incorporated and IGI, University of California at Berkeley.

## Author Contributions

JJ and M-JC designed the experiments. DD and MG constructed vectors, and JJ, EZ, DT and DR performed the experiments. BS and M-JC supervised the experiments. JJ, EZ, DT and M-JC contributed towards writing the manuscript. CG, J-PM, DL and RS provided protocol information, technical training and donor materials. All authors read and approved the manuscript.

## Abbreviations

DTT: Dithiothreitol
ED3: Embryo development 3% sucrose
ED4: Embryo development 4% sucrose
ED6: Embryo development 6% sucrose
EDL: Embryo development light
FPE: Fluorescent protein expression
nptII: Neomycin phosphotranferase II
OD600: Optical density at 600nm
PCG: Primary callus growth
QSE: Quaternary somatic embryo
SCG: Secondary callus growth
T-DNA: Transfer-DNA
TSE: Tertiary somatic embryo
YFP: Yellow fluorescent protein

