## Supplementary material for "Screening of Cultivars for Tissue Culture Response and Establishment of Genetic Transformation in a High-yielding and Disease-resistant Cultivar of *Theobroma cacao*": Jones et al_Supplemental_Fig1.pdf

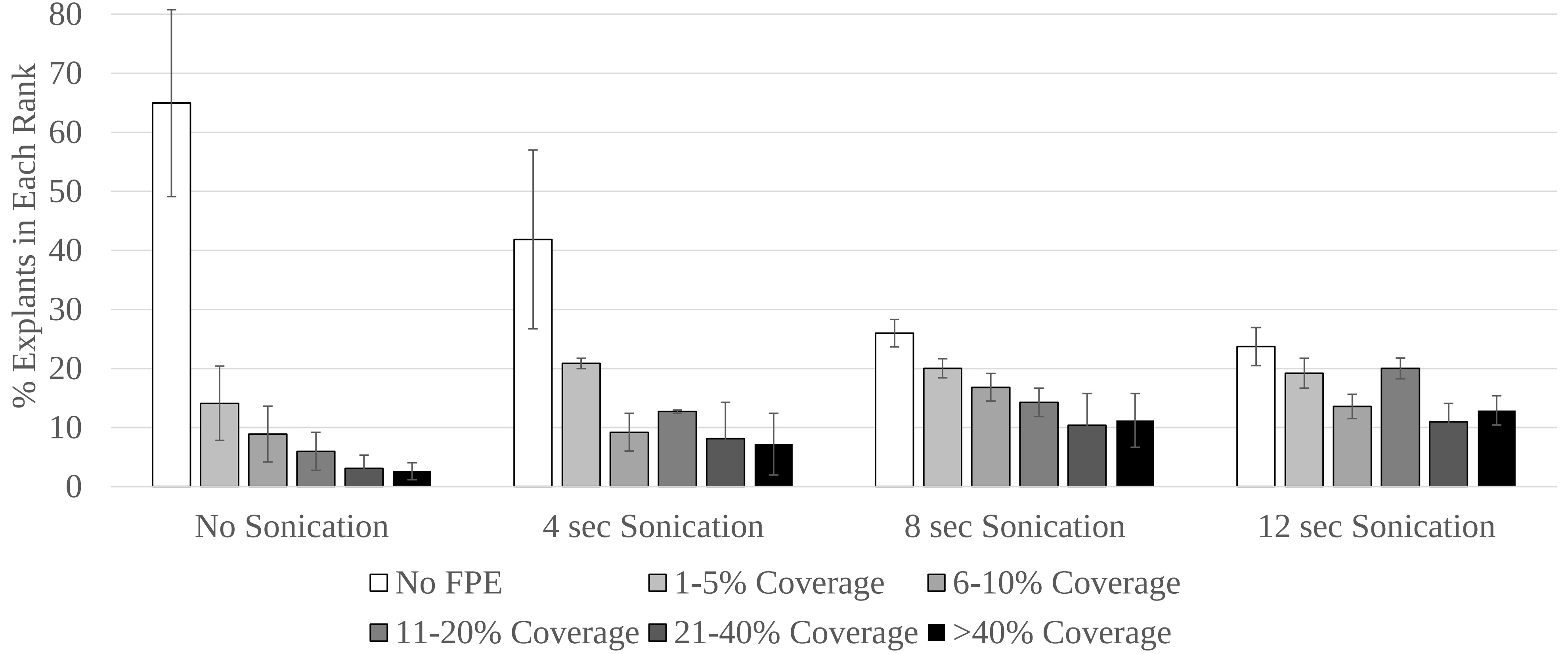


**Fig. S1** Effect of sonication treatment on transient FPE coverage in INIAPG-038. Cotyledon explants were observed in six ranks based on FPE coverage: No FPE, 1-5% FPE coverage, 6-10% FPE coverage, 11-20% FPE coverage, 21-40% FPE coverage and greater than 40% FPE coverage. Each histogram represents the mean level (±standard error) from between 2 and 3 replicates
